# Isotope tracing-based metabolite identification for mass spectrometry metabolomics

**DOI:** 10.1101/2025.04.07.647691

**Authors:** Deniz Secilmis, Arjana Begzati, Nina Grankvist, Irena Roci, Jeramie Watrous, Amit R. Majithia, Gordon I. Smith, Samuel Klein, Mohit Jain, Roland Nilsson

## Abstract

Modern mass spectrometry-based metabolomics is a key technology for biomedicine, enabling discovery and quantification of a wide array of biomolecules critical for human physiology. Yet, only a fraction of human metabolites have been structurally determined, and the majority of features in typical metabolomics data remain unknown. To date, metabolite identification relies largely on comparing MS^2^ fragmentation patterns against known standards, related compounds or predicted spectra. Here, we propose an orthogonal approach to identification of endogenous metabolites, based on mass isotopomer distributions (MIDs) measured in an isotope-labeled reference material. We introduce a computational measure of pairwise distance between metabolite MIDs that allows identifying novel metabolites by their similarity to previously known peaks. Using cell material labeled with 20 individual ^13^C tracers, this method identified 62% of all unknown peaks, including previously never seen metabolites. Importantly, MID-based identification is highly complementary to MS^2^-based methods in that MIDs reflect the biochemical origin of metabolites, and therefore also yields insight into their synthesis pathways, while MS^2^ spectra mainly reflect structural features. Accordingly, our method performed best for small molecules, while MS^2^-based identification was stronger on lipids and complex natural products. Among the metabolites discovered was trimethylglycyl-lysine, a novel amino acid derivative that is altered in human muscle tissue after intensive lifestyle treatment. MID-based annotation using isotope-labeled reference materials enables identification of novel endogenous metabolites, extending the reach of mass spectrometry-based metabolomics.

## Introduction

Mass spectrometry-based metabolomics is a key technology for monitoring human physiology, providing information on nutrients, endogenous metabolites, signaling molecules, environmental exposures, and other biological factors. Large-scale studies of metabolites in human populations have identified drug targets and uncovered biomarkers of disease pathogenesis and drug response ^1–3^. Yet, the human metabolome is vast and chemically diverse, and most human metabolites still remain to be discovered ^4^. While modern mass spectrometers can detect tens of thousands of peaks in a single sample ^5,6^, identifying the metabolites that give rise to these peaks remains a critical bottleneck in metabolomics. While transcriptomics and proteomics methods benefit from a reference genome against which observed sequences can be mapped, no such reference exists for metabolites. Instead, metabolite identification in mass spectrometry relies on measuring various physical characteristics of a metabolite which can be compared against known compounds ^7^, and the confidence in each metabolite identification depends on the specificity of those characteristics. Ion mass/charge ratio (m/z) and chromatographic retention time are easy to measure, but have limited specificity since other compounds may have identical m/z (isomers) and similar retention time ^8^. Fragmentation spectra from tandem mass spectrometry (MS^2^) can sometimes provide high specificity, but cannot always discriminate between related compounds ^9^, and high-quality MS^2^ spectra can be difficult to obtain for less abundant molecules in complex samples. Ion mobility, which measures the cross-section area of ions ^10^, provides complementary data that helps increase specificity, particularly for lipids and other large molecules. In all cases, a known compound is required to compare against, prompting the development of databases of MS^2^ spectra ^11,12^ and ion mobility data ^13^ for large numbers of reference compounds. While powerful for annotating unknown peaks arising from already known metabolites (“known unknowns”), this strategy cannot discover truly novel compounds (“unknown unknowns”) for which no reference compound exists, posing a fundamental limitation for metabolite discovery ^14^.

To enable discovery of novel metabolites, two main approaches have been explored. One strategy is to computationally generate collections of hypothetical chemical structures ^15,16^ and predict their MS^2^ fragmentation patterns ^17,18^ to generate libraries of “synthetic” reference spectra against which unknown peaks can be matched. A second, orthogonal approach is to quantify the similarity between MS^2^ fragmentation spectra of distinct but related compounds, which can give clues to the structure of an unknown metabolite of interest ^19,20^. Both approaches can enable discovery of novel metabolites ^19,21^, provided that high quality, informative MS^2^ spectra can be obtained. Yet, despite these advances, metabolite identification remains a challenging problem, and the majority of metabolites in typical non-targeted mass spectrometry datasets are still unknown.

Stable isotope incorporation is an alternative, less explored characteristic that could potentially be exploited for metabolite identification. When cells or tissues are provided nutrients containing stable isotopes such as carbon-13 (^13^C), those isotopes will be incorporated into metabolic products at specific positions, forming isotope patterns that reflect the biochemical synthesis pathway used. These isotope patterns can be highly specific, suggesting that they could be used to identify metabolites of interest. Importantly, isotope incorporation reflects the biochemical origin of a metabolite, and is therefore orthogonal to MS^2^ data which reflects structural features. For example, the isomers ribonate and xylonate are difficult to distinguish by MS^2^ as they are structurally similar, but could readily be identified from ^13^C tracing data since their biosynthesis pathways differ ^22^. We therefore reasoned that it might be possible to systematically identify endogenous metabolites in nontargeted mass spectrometry data based on a suitable isotope-labeled reference material. While ^13^C incorporation has been exploited to distinguish endogenous from non-endogenous metabolites in mass spectrometry ^23,24^, there is to our knowledge no methodology for systematic metabolite identification from isotope information.

Here, we introduce an approach and computational framework for isotope-based identification of unknown molecules in nontargeted mass spectrometry data. Using a reference material from human cells labeled with multiple ^13^C substrates, this approach allowed us to annotate hundreds of endogenous metabolites. As an illustrating example, we report the discovery of trimethylglycyl-lysine (TMGL), a previously unknown human metabolite.

## Results

### Metabolite identification using mass isotopomer distributions

Since biochemically related metabolites should exhibit similar patterns of isotope incorporation, we reasoned that unknown metabolites could be identified by comparing these patterns against known compounds. In mass spectrometry data, incorporation of ^13^C into an *n*-carbon metabolite is reflected by its mass isotopomer distribution (MID), defined as the fractional abundance of its *n* + 1 mass isotopomers (MIs) ^13^C_0,_ ^13^C_1, …,_ ^13^C_n_. In the simplest case where two metabolites are related via a biochemical pathway A → B that preserves the carbon skeleton, their MIDs should be virtually identical, and their pairwise distance very small. For example, in human epithelial cells cultured with U-^13^C_6_- glucose, we observed an unknown metabolite with predicted formula C_6_H_6_O_6_ whose MID was very close to that of citrate (Fig 1a), strongly suggesting that the unknown metabolite is aconitate, which is formed from citrate (C_6_H_8_O_7_) by removing a water molecule. On the other hand, if two metabolites A and B are related through a pathway A + C → B where C contains additional carbons, then the MID of B should theoretically be a convolution ^25^ of the MIDs of A and C (see Methods for details). Such convolutions can yield quite complex MIDs: for example, in the same U-^13^C_6_-glucose tracing experiment, convolution of the measured MIDs of UDP and glucose-6-phosphate (Fig 1b) resulted in an MID with prominent ^13^C_5_ and ^13^C_6_ MIs due to the ribose (C_5_) and glucose (C_6_) moieties, but also many less obvious MIs that arise from various combinations of MIs in UDP and glucose. Nevertheless, the predicted convolution agreed well with the measured MID of UDP-glucose (Fig 1b) in this experiment. This suggests that a general measure of biochemical relatedness between any two metabolites A and B could be defined by comparing the MID of B to the convolution A + C. However, a fundamental difficulty is that the metabolite C, which we refer to as the convolutant, is in general unknown. To solve this problem, we defined the MID distance between A and B to be the smallest Euclidean distance between the convolution A + C and the measured MID across all possible convolutants C in the experimental data (Fig 1c). Metabolites A, B for which this distance is small are then predicted to be biochemically related.

**Figure 1.**
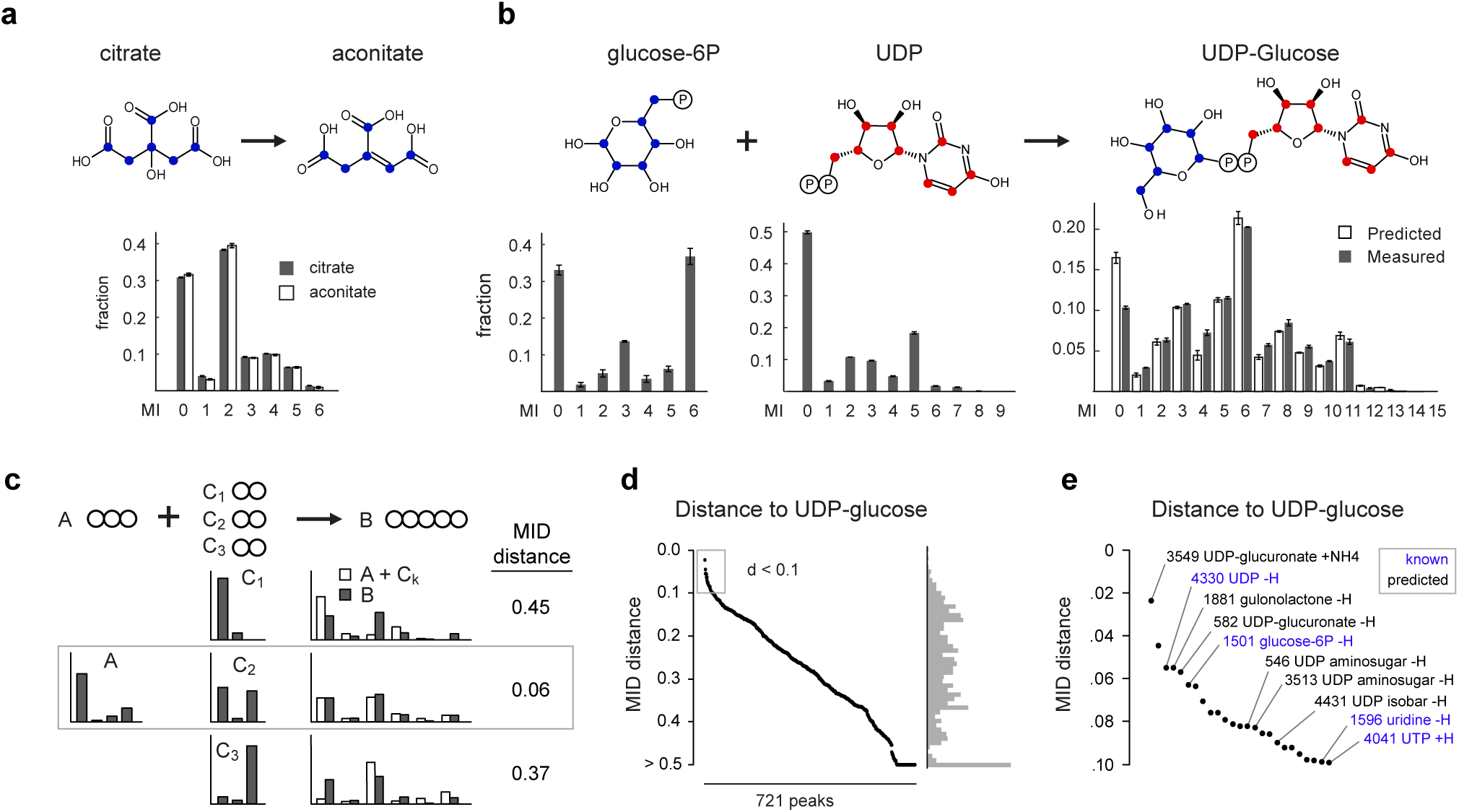
Principle of MID-based metabolite identification. **a**, Mass isotopomer distributions (MIDs) of citrate and aconitate in cell extracts. Blue dots indicate carbon atoms. MI, mass isotopomer. **b**, MIDs of glucose-6-phosphate, uridine-diphosphate (UDP) and UDP-glucose in cells, as well as predicted convolution MID (left). Red and blue dots indicate transferred carbon atoms. **c**, schematic for computation of the MID distance between hypothetical molecules A and B with convolutants C_1_, C_2_ and C_3._ Circles indicate carbon atoms; bar graphs indicate MIDs. The MID distance between A and B equals the Euclidean distance between the best matching convolution A + C_2_ and B (middle row). **d**, MID distances between UDP-glucose and 721 putative metabolites. **e**, zoom in on the region indicated in gray in (d), with predicted (black) and previously known (blue) compounds indicated. Error bars in (a, b) indicate standard deviation from n = 3 independent cultures.

To test if our MID distance could be used to identify metabolites in an unbiased fashion, we first performed an untargeted analysis of the U-^13^C-glucose labeling experiment. After filtering out mass spectrometry artefacts, we obtained 1,886 peaks representing potential metabolites. We further discarded peaks having low ^13^C enrichment or mass isotopomers colliding with other co-eluting peaks, resulting in a total of 721 peaks representing potential metabolites with varying degree of ^13^C incorporation (Extended Data Fig 1a, Suppl Table 1). We then searched for metabolites related to UDP-glucose by computing the MID distance between UDP-glucose and every other metabolite. Remarkably, the closest neighbors included UDP and glucose-6P, as well as UDP-related metabolites such as uridine (Fig 1d,e). Moreover, for the pair (UDP-glucose, glucose-6P), the best convolutant was UDP. The top neighbors of UDP-glucose also included unannotated peaks whose formula was consistent with other UDP-coupled aminosugars, leading to several new predicted metabolites. For peak 582, predicted to be UDP-glucuronate, we confirmed the identity by MS^2^ acquired from a pure standard (Extended Data Fig 1bc). These results demonstrate that it is possible in principle to identify unknown metabolites based on ^13^C MIDs.

### Multiple isotope labeling experiments increase specificity

Since human cell biomass is formed not only from glucose but also to a large extent from the proteinogenic amino acids ^26^, a single ^13^C substrate is expected to label only a subset of human metabolic products. To obtain better coverage and specificity, we generated a reference material of cell extracts from 20 parallel ^13^C labeling experiments, each substituting one amino for the U-^13^C-labeled form (Fig 2a, Extended Data Fig 2a). Each such experiment might label a different moiety of a given metabolite of interest, depending on its biochemical origin (Fig 2a). While glucose was the largest source of carbon, resulting in >5% ^13^C enrichment in more than half of all LCMS peaks, each ^13^C substrate contributed labeling to a particular group of peaks (Fig 2b). We detected labeling from more than one ^13^C substrate in 56% of all peaks, demonstrating synthesis from multiple nutrients. For example, glutathione exhibited a rich pattern of ^13^C labeling from cysteine, glucose, glutamine, glutamate, glycine and serine (Fig 2c), consistent with the known pathway of glutathione synthesis from these substrates. Hence, combining MIDs across labeling experiments should yield more informative patterns and higher specificity than can be obtained with any single experiment alone.

**Figure 2.**
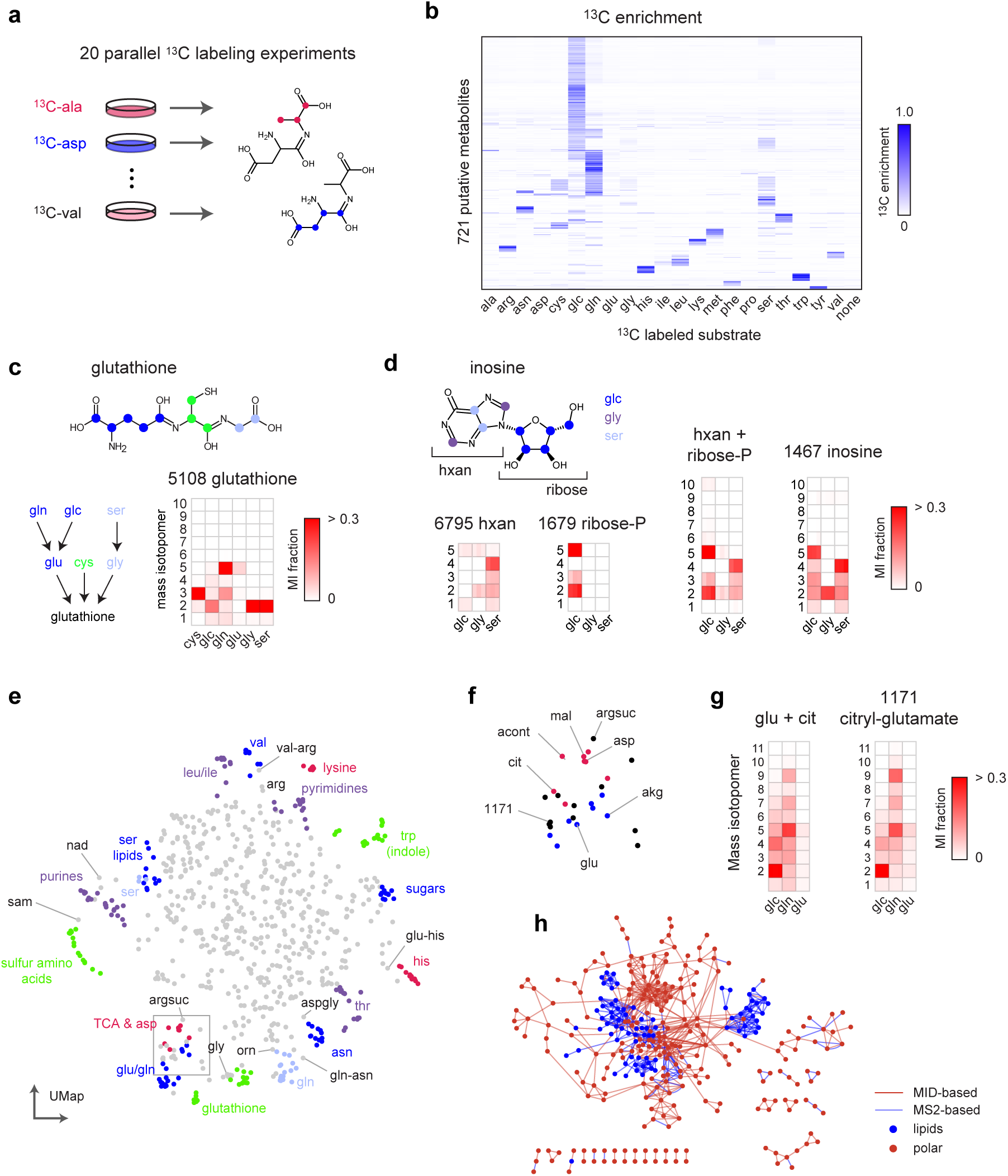
MID-based metabolite identification from multiple tracers. **a**, schematic of parallel isotope labeling experiments. Each experiment may label a different moiety of a compound of interest. **b**, Heatmap of ^13^C enrichment in 721 putative metabolites from ^13^C labeling experiments in cell extracts with indicated substrates. glc, glucose; control, unlabeled (^12^C) control culture. **c**, Glutathione structure (top), simplified scheme of glutathione synthesis (left), and MIDs of glutathione across labeling experiments (right), shown as a heat map of mass isotopomer (MI) fractions. Experiments with substantial ^13^C labeling are shown. **d**, Structure of inosine with origin of carbons indicated by color, and MIDs of hypoxantine (hxan), ribose-phosphate, their computed convolution, and inosine. **e**, UMap projection of all pairwise MID distances between 721 putative metabolites, with clusters of related metabolites highlighted. sam, S-adenosylmethionine; argsuc, argininosuccinate. **f**, zoom in on region of (e) with TCA cycle and glutamate-related metabolites indicated. **g**, MIDs of peak 1171 (citryl-glutamate) and the convolution of glutamate and citrate. **h**, Overlaid networks derived from MID-based distance (d < 0.7) and MS^2^-based molecular networking. Lipids (amphipathic) and polar compounds are indicated. z

We next generalized the MID distance measure to take advantage of multiple labeling experiments. Provided that all experiments interrogate the same metabolic state, two biochemically related compounds should have similar MIDs in every experiment. For example, the convolution of hypoxanthine and ribose-phosphate MIDs was similar to the MID of inosine (which is formed from these metabolites during purine salvage) in each experiment that resulted in ^13^C labeling (Fig 2d). Following this rationale, we took the sum of Euclidean distances between MIDs across all experiments, and then searched for the best convolutant minimizing this sum, as above (Fig 1c). We computed this MID distance between all pairs of peaks (Extended Data Fig 2b) and visualized the result using UMap projection. Reassuringly, this analysis grouped related metabolites into clusters reflecting their biochemical origin (Fig 2e, Suppl Table 2). For example, TCA cycle metabolites clustered together and were located near glutamate-related metabolites (Fig 2f), reflecting their close relationship via alpha-ketoglutarate, while serine-derived phospholipids were located near serine (Fig 2e). It should be kept in mind that, as complex biochemical relationships between metabolites cannot be perfectly represented in 2D, the UMap projection provides only an approximation of the actual MID distance (Extended Data Fig 2c). For example, arginine was located near valine in the UMap due to MID similarity to an arginyl-valine dipeptide, while the arginine metabolite argininosuccinate was placed near TCA cycle metabolites due to its succinate moiety (Fig 2e). Yet, argininosuccinate was the third closest neighbor of arginine in the actual MID distance. Interestingly, metabolites tended to be located in-between their biosynthetic precursors in the UMap projection. For example, the methyl donor S-adenosylmethionine was positioned between methionine metabolites and purines (Fig 2e), reflecting that it contains both methionine and purine moieties. Similarly, we noticed one unannotated compound positioned near glutamate and citrate (Fig 2f), whose MID closely matched their convolution (Fig 2g). We predicted this compound to be β-citryl-glutamate, a little-known metabolite originally discovered in brain ^27^, and confirmed this prediction by retention time and MS^2^ of a pure standard (Extended Data Fig 2d,e). Taken together, these data show that MID distances based on parallel ^13^C experiments reflect biochemically related metabolites.

To compare the MID-based distance against methods based on structural similarity, we next performed MS^2^-based molecular networking ^20^ on the HMEC data set. This generated a total of 318 connections among 140 of the 721 metabolites (20%). We then compared this network against the network obtained by thresholding the pairwise MID distances at a conservative cutoff (< 0.7). We observed that the MS^2^ network mainly connected lipids and other large, structurally complex molecules, while the MID-based distance generated more connections between small metabolites (Fig 2h). This highlights the complementarity of these two approaches, where fragmentation-based methods appear more suitable for large molecules, while MID-based annotation better recovers the biochemical relationships between small, central metabolites.

### Evaluation of the MID distance measure on simulated data

We next sought to understand in more depth how MID distance reflects biochemical similarity. In realistic metabolic networks, metabolites can be formed via multiple pathways, resulting in complex MIDs. To assess the accuracy of MID-based distances in such a situation where the ground truth is known, we performed simulation studies (Figure 3a) using a large atom-level model of central human metabolism (Figure 3b) consisting of ∼500 known reactions and using 25 substrates to synthesize metabolic products ^28^. We simulated 25 time-course isotopic labeling experiments, each with a different ^13^C-labeled substrate. To reflect that isotopic labeling in batch cultures is usually incomplete, we sampled MIDs at a time point before attaining steady state (Extended Data Fig 3a). From this simulated data, we computed all pairwise MID distances for all 25 labeling experiments. The UMap projection of these distances generally organized metabolites into the expected metabolic pathways (Figure 3c, Suppl Table 3), similar to our results from cultured human cells (Figure 2e). To investigate more closely how the MID distance can capture known biochemistry, we performed a local UMap projection of the TCA cycle (Fig 3d). Remarkably, this generally placed reactions into the correct sequence, suggesting that MID distances can be used to discover the local structure of metabolic pathways.

**Figure 3.**
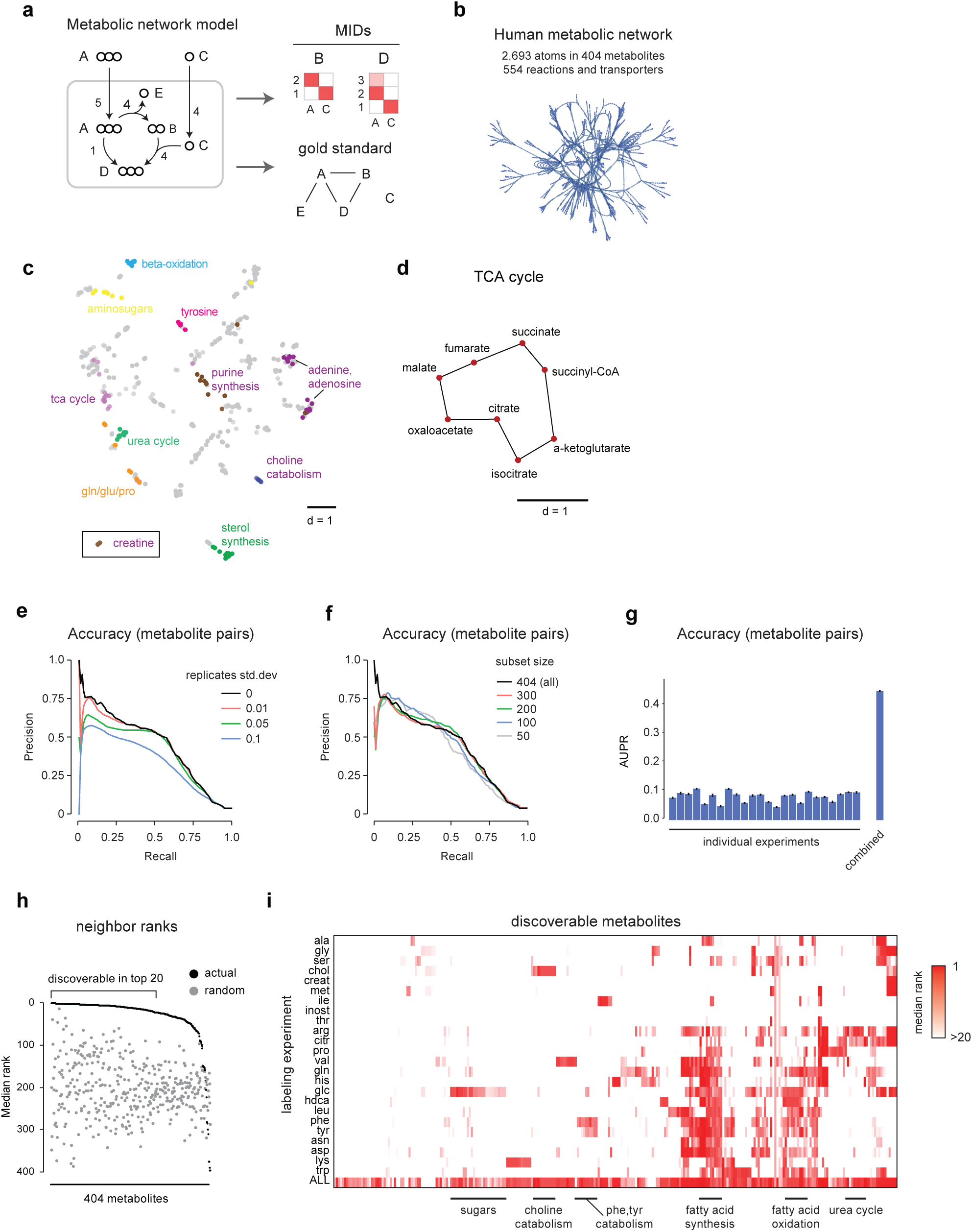
Computational evaluation of the MID distance. **a**, simple example of simulation study, where a metabolic network is used to simulate MIDs of metabolites B and D using labeled substrates A and C, and also to compute a “gold standard” for pairwise biochemical relatedness (see Methods). **b**, atom-level connectivity structure and summary statistics for the human metabolite network model used. **c**, UMap projection of MID distances from simulated data, from all experiments combined. Groups of related metabolites are highlighted. The creatine cluster was far separated and is shown as an inset. **d**, Local UMap projections of the TCA cycle, with lines indicating known biochemical reactions. **e**, Precision-recall curves for accuracy of the MID distance, compared to the gold standard derived from the human metabolic network, at indicated sampling noise levels. **f**, Precision-recall curves as in (f) on random metabolite subsets, simulating incomplete measurements. **g**, Area under the precision-recall curve (AUPR) for MID distances computed from each individual experiment and from all experiments combined. **h**, median rank by MID distance of the true neighbors of each metabolite; gray, ranks of randomly chosen neighbors. **i**, clustered heat map of ranks as in (i), for MID distances computed from each individual experiment, and from all experiments combined. Selected metabolite groups are indicated.

To quantitatively assess the accuracy of the MID distance in capturing biochemical relations between metabolites, a “gold standard” measure of biochemical relatedness is required (Fig 3a). Conventional metabolic pathways are not suitable here, as their definition is subjective, and one metabolite can often be placed in multiple pathways. As a systematic, pairwise measure of biochemical relatedness, we computed the fraction *f* of carbon contributed from one metabolite to another (see Methods), which reflects the biosynthetic origin of each metabolite. We found that *f* captured known metabolic relationships well (Extended Data Fig 3b). Defining “biochemically related” as *f* > 1/2 served that the MID distance recovered about half of all such pairs at 50% precision, somewhat depending on the amount of added measurement noise (Fig 3e). Importantly, when a random fraction of the metabolites was dropped out, accuracy did not decline much on average (Fig 3f), indicating that the MID distance is effective also when only a subset of cellular metabolites are measured, which naturally is the case in practice. However, accuracy did vary markedly when taking small random subsets of the data (Extended Data Fig 3c), suggesting that the MID distance becomes inaccurate if “key” metabolites happen to be missing. We also wondered if comparable accuracy could be obtained using fewer isotope tracing experiments, which might simplify the generation of labeled reference material. However, no single experiment achieved the accuracy obtained using all experiments (Fig 3g), and when iteratively adding the next-best experiment, about half of the experiments were needed to achieve comparable accuracy (Extended Data Fig 3d). While these results depend on particulars of the underlying simulation, they indicate that using multiple ^13^C tracers is essential for good accuracy on human metabolites.

To evaluate the ability of the MID distance to identify new compounds, we assessed how often biochemically related metabolites were found among the closest neighbors of each metabolite. Using the MID distance based on all simulated experiments, we find that more than half of all metabolites were “discoverable”, defined as recovering at least half of the true biochemically related metabolites in the top 20 (Fig 3h). When assessing accuracy using individual experiments, we found that different tracers enable discovery in different metabolic pathways, as might be expected (Fig 3i). These results indicate that, in principle, MIDs capture enough information to allow identifying a large fraction of unknown metabolites in a human metabolic network with good accuracy.

### Systematic evaluation of MID-based metabolite identification

To systematically evaluate the effectiveness of MID-based metabolite annotation in practice, we next annotated all 721 peaks in the HMEC data set (Fig 4a). We started from a set of 101 known peaks whose identity was established beforehand based on retention time and MS^2^ spectra matching against pure standards (Suppl Table 1). We excluded 183 peaks that were considered likely phospholipids, which all exhibited similar MIDs typical of *de novo* synthesis of fatty acyl chains (Extended Data Fig 4a). For the remaining 437 peaks, we first retrieved candidate annotations by matching on m/z against HMDB, resulting in an average of 8 candidates for each peak (Extended Data Fig 4b). We then determined for each peak the closest known metabolites by MID distance, and selected the candidate compound that was most similar to those known neighbors. For example, peak 4889 with predicted formula [C_12_H_22_N_3_O_8_]^+^ matched four candidate compounds in HMDB, of which aspartyl-glucosamine was predicted to be the true compound since the closest neighbors of this peak included asparagine and acetyl-glucosamine (Fig 4b). In this manner, we could predict an identity for 272 of the 437 unknown non-lipid peaks (62%) based on MID distance, representing 146 unique structures (Fig 4c; Suppl Table 1). To validate these predictions, we obtained pure standards for aspartyl-glucosamine and three other predicted compounds (Extended Data Fig 4c–e), and in all cases confirmed their identity by retention time and MS^2^ fragmentation pattern (Extended Data Fig 4f–i). For comparison, MS^2^ matching using GNPS predicted identities for 122 of the non-lipid peaks (28%), comprising 100 unique structures (Fig 4c; Suppl Table 4). However, MS^2^ matching also gave predictions for 55 of the lipid peaks (31%), again underscoring the complementary strengths of these approaches.

**Figure 4.**
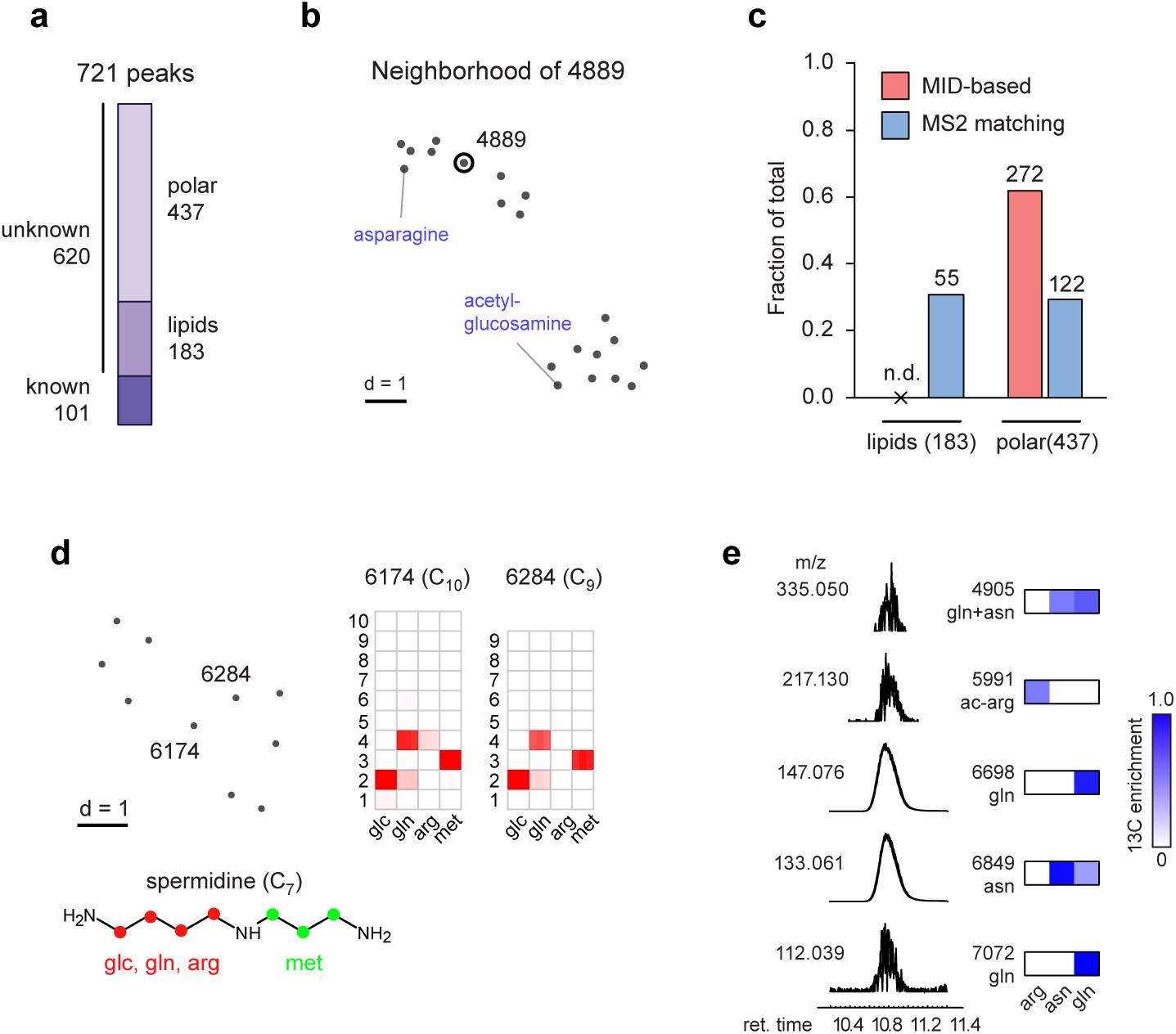
Systematic evaluation of MID-based metabolite identification. **a**, Peaks in the HMEC data categorized as previously known; unknown lipids, and unknown polar compounds. **b**, UMap projection of the neighbors of peak 4889 (circled) according to the MID distance. Nearby known compounds indicated in blue. **c**, number and fraction of unknown lipids and polar metabolites determined by MID-based and MS^2^-based metabolite identification. n.d, not determined. **d**, UMap projection and MIDs of unknown 6174 and 6284, consistent with a spermidine-related structure (bottom). **e,** chromatograms of peaks coeluting with glutamine (relative intensity), with m/z (left) and ^13^C enrichment from relevant labeling experiments shown as heatmaps.

For peaks where we could not predict a likely compound structure, partial information on structure and biochemical origin could often be obtained, and false positive annotations ruled out. For example, for peak 5872 with formula [C_9_H_14_N_3_O_4_]^+^, the closest MID neighbors suggested a histidine-derived compound formed by convolution with a 3-carbon intermediate (Extended Data Fig 4j), while none of the 5 candidates in HMDB were structurally related to histidine. We also noticed two related compounds whose MIDs were consistent with a spermidine moiety, but whose exact identity remains unclear (Fig 4d). We identified 97 peaks as in-source fragments or adducts by grouping coeluting peaks that were similar by MID distance. However, we also found examples of co-eluting peaks with markedly different MIDs, indicating distinct metabolites that are not resolved by chromatography. For example, glutamine co-eluted with several other species, of which some were likely glutamine fragments, but others exhibited distinct ^13^C labeling, ruling out glutamine as a source (Fig 4e). Hence, MID information helps recover metabolites that otherwise might have been mistaken for artefacts.

While metabolites that incorporate ^13^C are clearly formed from the labeled substrates, it remains possible that some products are formed spontaneously. For example, we predicted several glucosylated and lactoylated amino acids such as glucosyl-serine (Extended Data Fig 4k) and lactoyl-asparagine (Extended Data Fig 4l). These could be due to nonenzymatic addition of the reactive aldehydes glucose and methylglyoxal, respectively; however, since the glucosyl and lactoyl moieties were ^13^C-labeled, it seems likely that these reactions occurred in cells. N-lactoyl-amino may be formed by reverse proteolysis ^29^, and one predicted metabolite, lactoyl-phenylalanine (peak 1635; Suppl Table 1), has recently been reported to function as an endogenous appetite suppressor ^30^, raising the possibility that related metabolites have metabolic functions as well.

### Discovery of a new human metabolite

Among peaks lacking plausible candidates in HMDB, we noticed one unknown compound with ^13^C labeling from glycine, lysine, methionine and serine. This compound was closely related by MID distance to trimethyllysine and glycine (Fig 5a), and its MID indicated that it contained lysine, glycine and three methionine-derived methyl groups (Fig 5b). We first considered trimethyllysyl-glycine or glycyl-trimethyllysine as possible structures, but MS^2^ spectra of these compounds did not match that of the unknown compound, and in particular failed to explain a prominent MS^2^ fragment at m/z 100.076 (Extended Data Fig 5a,b). However, a trimethylglycyl-lysine (TMGL) structure (Fig 5c) explained all observed MS^2^ fragments (Extended Data Fig 5c), leading us to hypothesize that the unknown compound is TMGL. To test this hypothesis, we synthesized pure TMGL and found that its MS^2^ spectra and retention time closely matched that from HMEC extracts (Extended Data Fig 5c,d). As additional validation, we performed MS^2^ analysis of TMGL from cells labeled with either U-^13^C-lysine, U-^13^C-methionine, or with “deep labeling” medium ^22^ where all amino acids were fully ^13^C-labeled. In each case, we observed ^13^C mass shifts in MS^2^ fragments consistent with the TMGL structure (Fig 5d, Extended Data Fig 5e,f), and in particular confirmed that the methyl groups are located on the glycine amine group (Fig 5d). We therefore conclude that the unknown metabolite is indeed TMGL.

**Figure 5.**
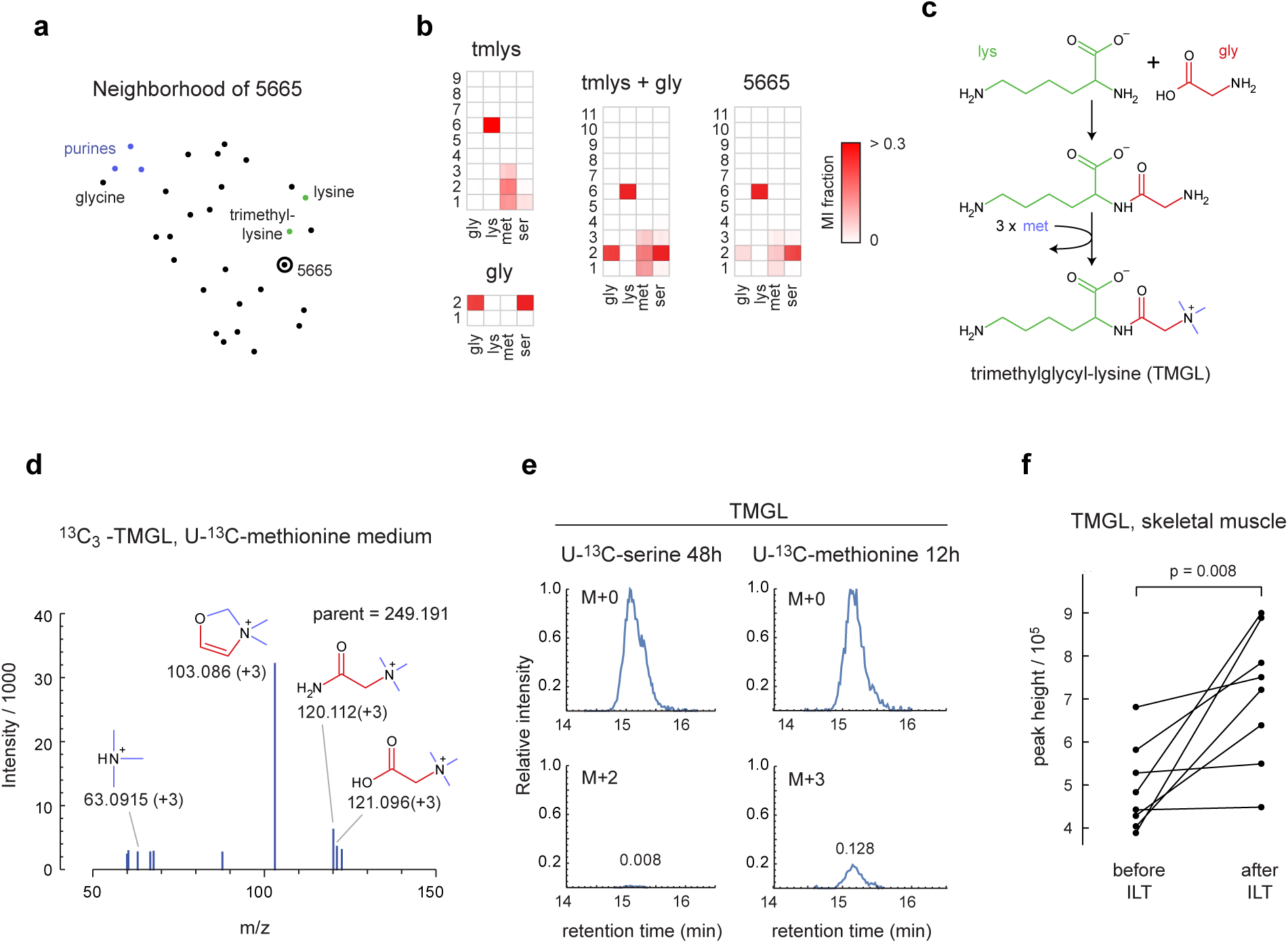
Discovery of the human metabolite trimethylglycyl-lysine (TMGL). **a**, UMap projection of the neighbors of peak 5665. **b**, MIDs of trimethyllysine (tmlys), glycine (gly), their convolution (tmlys + gly) and peak 5665. Experiments with substantial ^13^C labeling are shown. **c**, Putative synthesis pathway of TMGL from lysine, glycine and methionine. **d**, MS2 spectra of TMGL peak from cells labeled with ^13^C-methionine, showing expected ^13^C labeling of fragments containing methyl groups. **e**, Relative abundance of indicated TMGL indicated mass isotopomers, normalized to M+0 apex, in cells labeled with U-^13^C-serine or U-^13^C-methionine for indicated time periods. Numbers indicate MI fraction. **g**, Relative abundance of TMGL in human skeletal muscle before and after intensive lifestyle therapy (ILT).

We next wondered how TMGL might be synthesized in cells. While we observed ^13^C labeling of the TMGL methyl groups from ^13^C-methionine, free trimethylglycine (TMG; also known as betaine) was unlabeled (Extended Data Fig 6a), consistent with the established pathway where TMG is formed from choline ^31^. Also, in tracing experiments with methyl-^13^C-choline, TMG acquired ^13^C labeling but TMGL did not (Extended Data Fig 6b), indicating that TMGL is not derived from TMG. To probe the synthesis pathway, we performed transient ^13^C tracing experiments with U-^13^C-serine and U-^13^C-methionine, and observed that the glycyl moiety of TMGL labeled very slowly from U-^13^C-serine, while U-^13^C-methionine rapidly labeled the TMGL methyl groups (Fig 5e). This suggests that a pathway where the glycyl-lysine dipeptide is formed first by a slow process, perhaps as a byproduct of protein catabolism, followed by rapid trimethylation of the glycine amine group by a currently unknown enzyme (Fig 5c).

To our knowledge, this is the first report of the existence of TMGL in any organism. Its biological role is unknown. We observed that TMGL accumulated in spent medium from cell cultures (Extended Data Fig 6c), suggesting that it is released from cells, and therefore might also be excreted into the circulation. Considering that TMGL might arise from glycyl-lysine peptides derived from protein catabolism, we decided to investigate this metabolite in a physiological context where protein catabolism is increased. To this end, we analyzed TMGL in skeletal muscle from people with obesity and type 2 diabetes randomized to receive an 8-month intensive lifestyle therapy (ILT) that involved both dietary energy restriction and supervised exercise training, or a control group receiving standard care ^32^. We observed that TMGL was present in both plasma and muscle tissue (Extended Data Fig 6d,e). Muscle TMGL content increased markedly in the ILT group (Fig 5f), concurrent with marked weight loss (–16.6 ± 2.5%) and muscle remodeling ^32^, but not in the control group (Extended Data Fig 6f) where weight loss was minimal (–1.1 ± 1.0%). These data suggest a potential role for TMGL in human muscle physiology.

## Discussion

Our results demonstrate that stable isotope patterns can be leveraged to systematically annotate unknown LCMS peaks and discover novel compounds, including human metabolites of biomedical interest such as TMGL. While in this study metabolite identification relied solely on MIDs, isotope-based methods can easily be combined with MS^2^ spectra, ion mobility values and other characteristics. Given the complementary strengths of isotope-based and current fragmentation-based methods, we expect that such an integrated approach would outperform either method alone. Moreover, MS^2^ spectra from isotope-labeled samples can provide more detailed information on positional labeling ^33^, which could be exploited to further increase precision of isotope-based identification.

Importantly, the MID-based approach does not only identify metabolites, but also helps uncover the reactions or pathways that give rise those metabolites. In the TMGL example, MID data predicts a novel pathway where a hitherto unknown enzyme (or enzymes) methylates the glycine nitrogen, likely using S-adenosylmethionine as methyl donor, since ^13^C-methionine labels the TMGL methyl groups. Moreover, our simulation studies indicate that the MID distance can recover a substantial fraction of biochemically related metabolite pairs, and this is corroborated by cluster analysis of cell extract data. However, such metabolite pairs may be separated by multiple enzymatic steps, and candidate relationships identified by MID distance should be carefully examined to determine plausible underlying biochemical reactions.

Several caveats should be pointed out. First, although two metabolites A and B separated by a large MID distance are likely unrelated, a small MID distance does not always mean that A and B are related, since there can exist convolutants C such that convolution A + C is similar to B simply by chance. In our experience, this tends to occur when A and B are molecules of very different size, or when isotopic enrichment is low. Such false positives could be reduced by restricting convolutants to a predetermined set of biochemical intermediates (methylation, acetylation, etc), but doing so can also lead to discarding truly related pairs when pathway intermediates are missing from the data. Isobaric compounds with very similar MIDs, such as different hexose moieties, are also difficult to distinguish and should be treated as uncertain. We also caution that our MID distance is not a true metric as it violates the triangle inequality, which may lead to unexpected results with clustering and projection methods such as the UMap. Other distance measures that better fit the information geometry of probability distributions ^34^ may be worth exploring here. Finally, “false” mass isotopomers caused by overlapping LCMS peaks are common in untargeted ^13^C mass spectrometry data, and further development of preprocessing methods, such as more powerful co-elution tests, will be important to achieve a streamlined workflow.

Although much remains to be explored, the MID-based approach adds a new, orthogonal technique to the toolbox of metabolite identification methods, which currently relies heavily on structural information. When integrated into existing workflows, we anticipate that this approach will markedly improve our ability to identify metabolites from untargeted data with high confidence. Future work includes the development of an isotope-labeled reference material to make the methodology broadly accessible to the biotechnology community. Being internal ratios, MIDs are highly reproducible across instrument types and laboratories, facilitating data sharing. We anticipate that isotope-based ID will be particularly important in biomedicine, where it can accelerate discovery of endogenous human metabolites and their biochemical origin.

## Supporting information

Data set 1

## Acknowledgements

This work was supported by Swedish Research Council Grant no. 2020-01631 to DS, NG and RN; SSF grant no FFL-12-0220 to NG, IR and RN; NIH grants P30 DK056341 (Washington University Nutrition and Obesity Research Center) and UL1 TR000448 (Washington University Institute of Clinical and Translational Sciences) to SK; and NIH grants R01DK129840 and R01HL159760 to AB and ARM. The authors would like to acknowledge Yaroslav Lyutvinskiy for assistance with mass spectrometry data processing.

## Methods

### Cell culture

Human Mammary Epithelial Cells (HMECs) immortalized with the human telomerase gene were kindly provided by Dr. William C. Hahn (Dana-Farber Cancer Institute, Boston, USA) and have been previously described ^35^. HMECs were grown in custom-synthesized Mammary Epithelial Basal Medium (MCDB) 170 according to the original formulation ^36^, supplemented with 1% Mammary Epithelial Growth Supplement (MEGS) (S0155, Gibco), 100 units/ml penicillin and 100 μg/ml streptomycin (15140122, Thermo Fisher Scientific). Cells were maintained in a humidified atmosphere of 5% CO2/95% air at 37°C and washed and detached using ReagentPack Subculture Reagents (CC-5034, Lonza). HCT-116 (ATCC, CCL-247) and MCF7 (ATCC, HTB22) cells were maintained in RPMI-1640 medium supplemented with 5% dialyzed FBS (Gibco, 16140–071), prepared by dialysis in 10K MWCO dialysis tubing (Thermo Fisher Scientific #88245). All experiments were performed in triplicates.

### Isotope tracing

For isotope tracing in HMECs, 400,000 cells were seeded at day 0 in a T25 flask containing 5mL medium, and incubated overnight. On day 1, the medium was changed to an isotope tracing medium of the same molar composition, but with one nutrient exchanged for the corresponding U-^13^C-labeled substrate (Cambridge Isotope Laboratories). On day 2, each T25 flask culture was detached and seeded into two T25 flasks containing the same medium. On day 4, cells were detached and seeded into 6-well plates at 250,000 cells/well in 2mL of medium, in triplicate for each tracer. On day 7 (after roughly 6 cell divisions), each multi-well plate was placed on ice, medium was aspirated and cells were washed twice with 1 mL of cold PBS. Then, 1 mL cold (–80°C) methanol (JT Baker, BAKR8402.2500, VWR) was added to each well, cells were scraped using a 17mm cell scraper (83.1830, Sarstedt), and cell extracts were carefully transferred to a new tube, vortexed for 30 seconds to break up aggregated cell material, and stored in −80°C until analysis. The isotope tracing experiment with methyl-^13^C_3_-choline in MCF7 cells has been previously described ^31^. For transient isotope tracing experiments with U-^13^C-serine and U-^13^C-methionine, HCT116 cells were seeded in 6-wells at a density calculated to yield 500,000 cells per well at extraction time, precultured overnight in unlabeled custom-synthesized RPMI-1640, switched to corresponding medium containing the respective ^13^C tracer, and extracted in the same way as above at the indicated time points.

### Human study

The data reported here are analysis of plasma and skeletal muscle samples collected from participants that completed a study evaluating the effect of intensive lifestyle therapy (ILT) versus standard care on cardiometabolic function (ClinicalTrials.gov identifier NCT01977560) ^32^. Participants were included in this study if they had an adequate amount of skeletal muscle tissue available before and after weight loss to assess tissue LTMG levels. A total of 14 people with obesity and type 2 diabetes (age 52.4 ± 2.6 years; body mass index 37.0 ± 1.5 kg/m^2^) participated. All procedures were conducted in the Clinical Translational Research Unit (CTRU) at Washington University School of Medicine in St. Louis, MO. The study was approved by the Human Research Protection Office at Washington University School of Medicine in St. Louis, MO, and all the participants provided written informed consent before participation. Study inclusion criteria and behavioral interventions have been described previously ^32^. After an overnight fast, a blood sample was obtained from a catheter inserted into a radial artery and a vastus lateralis muscle tissue sample was obtained by percutaneous biopsy by using a Tilley Henkel forceps (Sontec Instruments, Inc., Centennial, CO).

### Mass spectrometry

LCMS analysis of HMEC, MCF7 and HCT116 cell extracts as well as pure standards was performed using pHILIC liquid chromatography coupled to a Thermo QExactive orbitrap mass spectrometer, as previously described ^22^. Human plasma samples were thawed on ice, and 40µl plasma was combined with 160µl of ice-cold methanol and vortexed. For muscle tissue samples, 20µl of ice-cold extraction solvent made of 80:20 methanol:water was added per milligram of tissue to account for mass variation, and samples were homogenized with ∼300mg 1mm zirconium beads using 3 x 10 second cycles at 6,400 Hz. All samples were then incubated at –20C° for 30 minutes to allow for proteins to precipitate, vortexed and centrifuged at 10,000 rpm for 10 min at 4°C. Supernatants of plasma samples were transferred to new tubes, dried down *in vacuo* using a vacuum concentrator, re-suspended in 25μL of 80:20 methanol:water, vortexed, sonicated for 2 minutes, and centrifuged at 10,000 rpm for 10 min at 4°C. Supernatants of plasma and tissue samples were then transferred to LC-HRMS vials containing 200μl glass inserts. 2μL of plasma supernatant and 1μL of tissue supernatant, which contained metabolic material from 50μg of tissue, was injected for LC-MS/MS based metabolomics analysis as previously described ^22^.

### LCMS data processing

For HMEC extracts, untargeted LCMS peak detection was performed on a triplicate of unlabeled cells from Thermo .raw data files using our in-house developed software “Isotrack”, available on request from the authors. This resulted in 4,743 and 3,198 LCMS peaks in positive and negative mode, respectively, that were reproducibly detected in all three replicates. Mass spectrometry artefacts, natural isotopomers, adducts and fragments were removed using the NetID R package ^6^, resulting in 1,886 peaks representing putative metabolites. Unlabeled metabolites were then removed by requiring at least 10% ^13^C enrichment from at least one labeled substrate, and the remaining peaks were curated manually to remove cases where mass isotopomer peaks collided with other peaks (“false” isotopomers), resulting in a final set of 721 LCMS peaks (Suppl Table 1). The full set of mass isotopomer distributions are available as Suppl Data 1.

Peak areas for all ^13^C mass isotopomer (MI) peaks of each metabolite ion were computed using the mzAccess data access framework ^37^ accessed from Mathematica v.11 (Wolfram Research), with peak coordinates obtained either from the untargeted analysis (HMEC cells) or from known m/z and retention times (all other experiments). MIDs were calculated by dividing the area of each MI peak by the sum of all MI peak areas. Remaining false isotopomers were filtered by setting to zero any MI except the base isotopomer that was higher than 0.03 in more than 10 experiments after correction for naturally occurring ^13^C.

### MID distance measure

The MID distance measure was calculated as follows. For each pair of metabolites *A* and *B* containing *m* and *n* carbons respectively (*m* < *n*) we consider all metabolites (“convolutants”) *C* in the data set having *n* − *m* carbons. For each convolutant *C*, we compute the convolution *x*^*A*^ × *x*^*C*^ of the MID vectors *x*^*A*^and *x*^*C*^ elementwise as

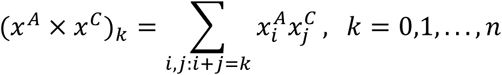

and then compute the MID distance *d*(*A*, *B*) as

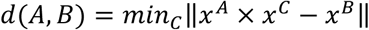

where ‖… ‖ is the Euclidean norm and the minimum is taken over all convolutants *C*. If no convolutant of size *n* − *m* was present in the data set, we set *d*(*A*, *B*) equal to the largest distance across all metabolite pairs. When using multiple MIDs from parallel tracing experiments, we minimize the sum of differences across all experiments *e*,

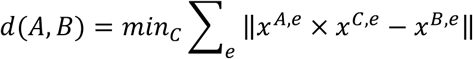

All distance calculations were performed on MID data without correction for natural ^13^C to avoid artefacts induced by the correction method. All MIDs shown were corrected for natural ^13^C prior to plotting.

### Metabolic network simulation

A previously described human metabolic network model ^28^ consisting of 554 reactions and 477 metabolites across four compartments was used for all simulations. A flux vector was chosen such that all reactions in the model was active at comparable flux magnitudes ^28^. The model was decomposed into elementary metabolite units (EMUs) and EMU reactions as previously described ^38^, and isotopic nonstationary simulation was done by generating time-dependent differential equations for each mass isotopomer of each EMU, which were numerically solved using Mathematica v11.0 (Wolfram Research, Inc). The simulation code is available at https://github.com/Nilsson-Lab-KI/nonstationary-mid-sim.

### Biochemical relatedness

The biochemical relatedness measure *f* was computed for each pair of metabolites A, B in the human metabolic network as follows. First, we removed all contributions from reactions producing A in the network so that A becomes a network substrate, and then fixed the isotopomer distribution of A to be fully labeled. We then simulated the steady-state MID of B by solving the steady-state EMU equations, at same flux state used for MID simulation. The ^13^C enrichment of B, defined for an n-carbon molecule as the sum 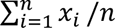, is then a measure of the fraction of carbon atoms in B derived from A at this flux state. Since the fraction of B derived from A does not necessarily equal the fraction of A derived from B, we define *f* to be the largest of the two values. Code for reproducing these computations is available at https://github.com/Nilsson-Lab-KI/nonstationary-mid-sim.

To compare the measure *f* against conventional metabolic pathways, we manually created a one-to-one mapping between metabolites in the human metabolic network model and conventional metabolic pathways. Metabolites considered to belong to more than one plausible pathway were left unassigned, resulting 278 metabolite-pathways mappings (Suppl Table 3).

### MID-based peak annotation

To identify candidate structures based on MIDs, we searched each unknown peak for previously known metabolites in its MID distance neighborhood. Potential candidates were generated by matching the unknown peak m/z against monoisotopic neutral mass of candidates obtained from HMDB within 10 ppm mass tolerance, considering M+H, M–H, M+NH_4_, M+H–H_2_O and M–H–H2O adducts. Predicted structures were generated by selecting candidates that (1) were structurally similar to nearby known peaks and (2) consistent with the observed number of ^13^C-labeled atoms from each tracer.

### MS^2^-based peak annotation

For peaks with unknown identity, we performed MS^2^ matching using the GNPS library search workflow ^12^. MS^2^ spectra were searched against publicly available spectral libraries, requiring that precursor and fragment m/z’s are within 0.2 Da, a minimum of 3 matched fragments, and a minimum cosine score of 0.8, considering the same set of adducts as for MID-based annotation (above). All hits to library spectra were manually reviewed to confirm accurate matches and to rule out false positives.

---

**Extended Data Figure 1.**
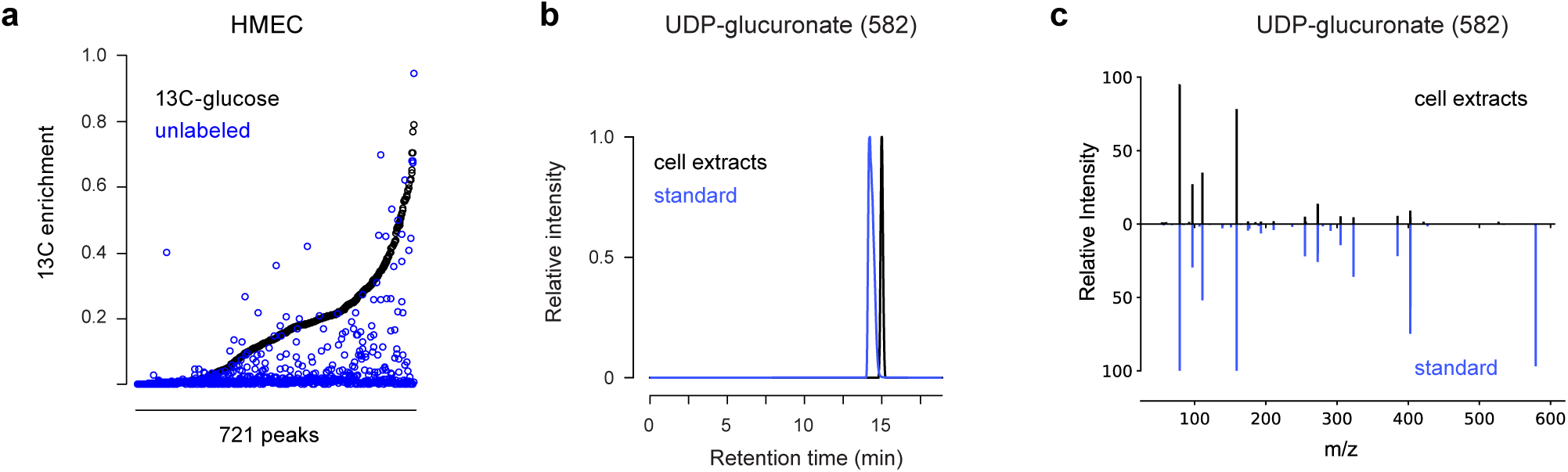
**a**, Distribution of ^13^C enrichment in 721 LCMS peaks in cells labeled with U-^13^C-glucose and in unlabeled cells. **b**, Chromatograms of peak 582 in cell extracts and a pure UDP-glucuronate standard. **c**, MS^2^ fragmentation spectra of peak 582 in cell extracts and a pure UDP-glucuronate standard.

**Extended Data Figure 2.**
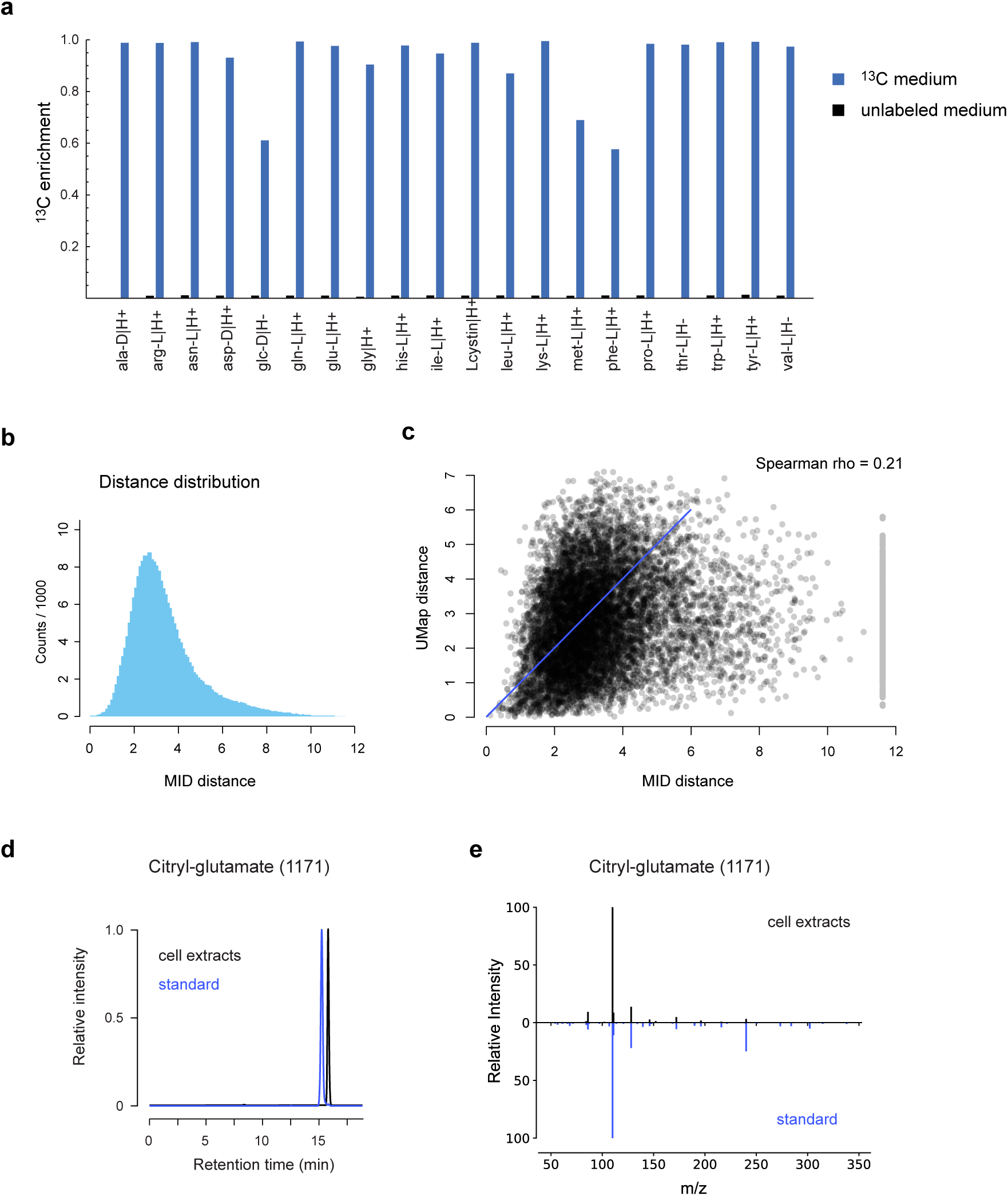
a, ^13^C enrichment of the indicated metabolites in the 20 corresponding U-^13^C-labeled media and in unlabeled medium. **b**, distribution of pairwise MID distances based on all tracers combined, corresponding to Fig 2e. **c**, scatter plot of MID distances vs. corresponding projected UMap distances. Line of identity indicated in blue. **d**, Chromatograms of peak 1171 in cell extracts and a pure citryl-glutamate standard. **e**, MS^2^ fragmentation spectra of peak 1171 in cell extracts and a pure citryl-glutamate standard.

**Extended Data Figure 3.**
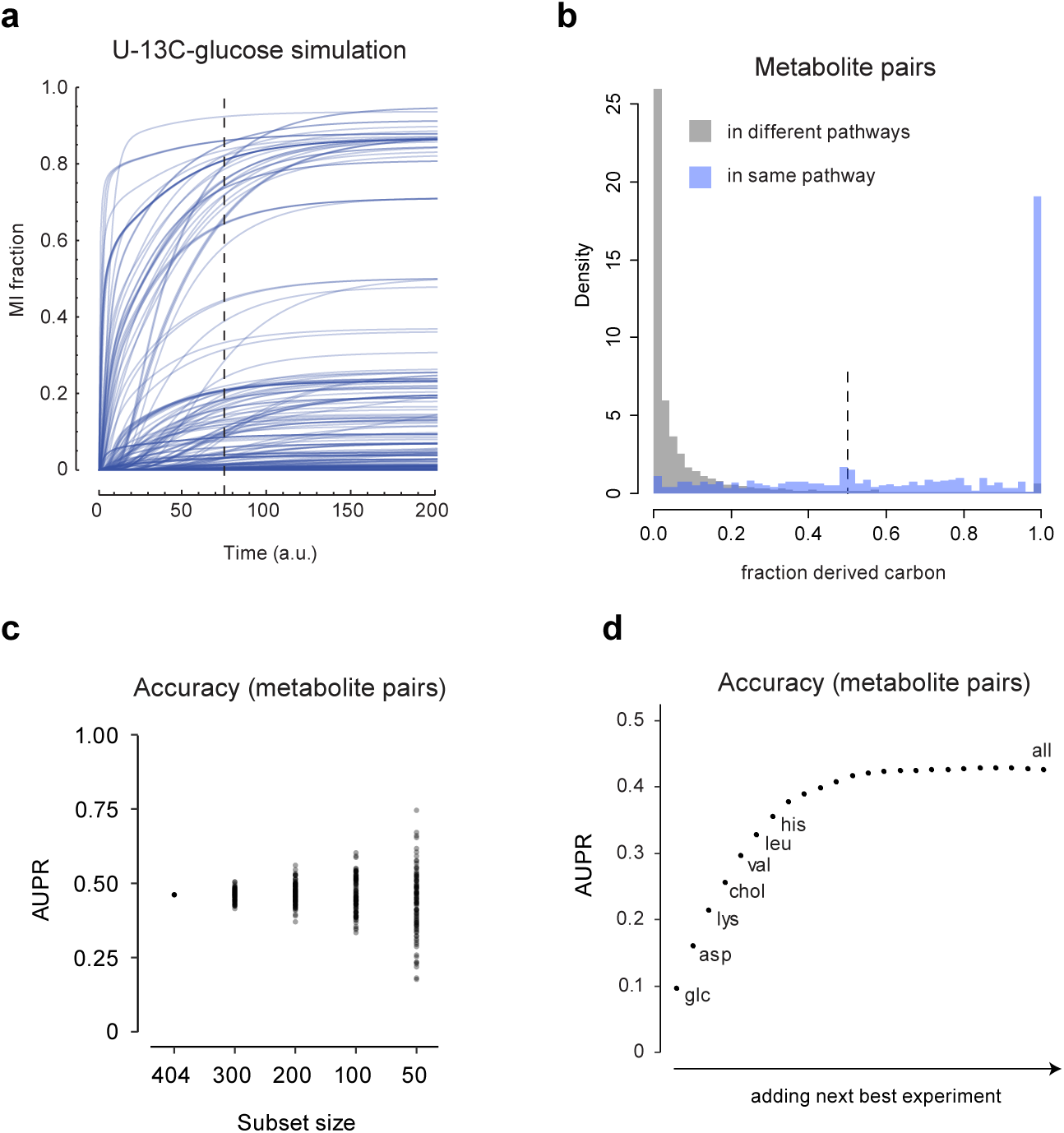
**a**, Time course of 3,097 mass isotopomer (MI) fractions in 404 simulated metabolites with U-^13^C-glucose as the labeled substrate. Dashed line indicates sampled time point. **b**, Histogram of the fraction *f* of derived carbons for all metabolite pairs, in either the same or different pathway. **c**, Variation in area under the precision-recall curve (AUPR) for MID distances computed from all experiments combined, for 100 random metabolite subsets of indicated sizes. **d**, AUPR of MID distances when iteratively adding next-best experiment. Experiments chosen in the first few iterations are indicated.

**Extended Data Figure 4.**
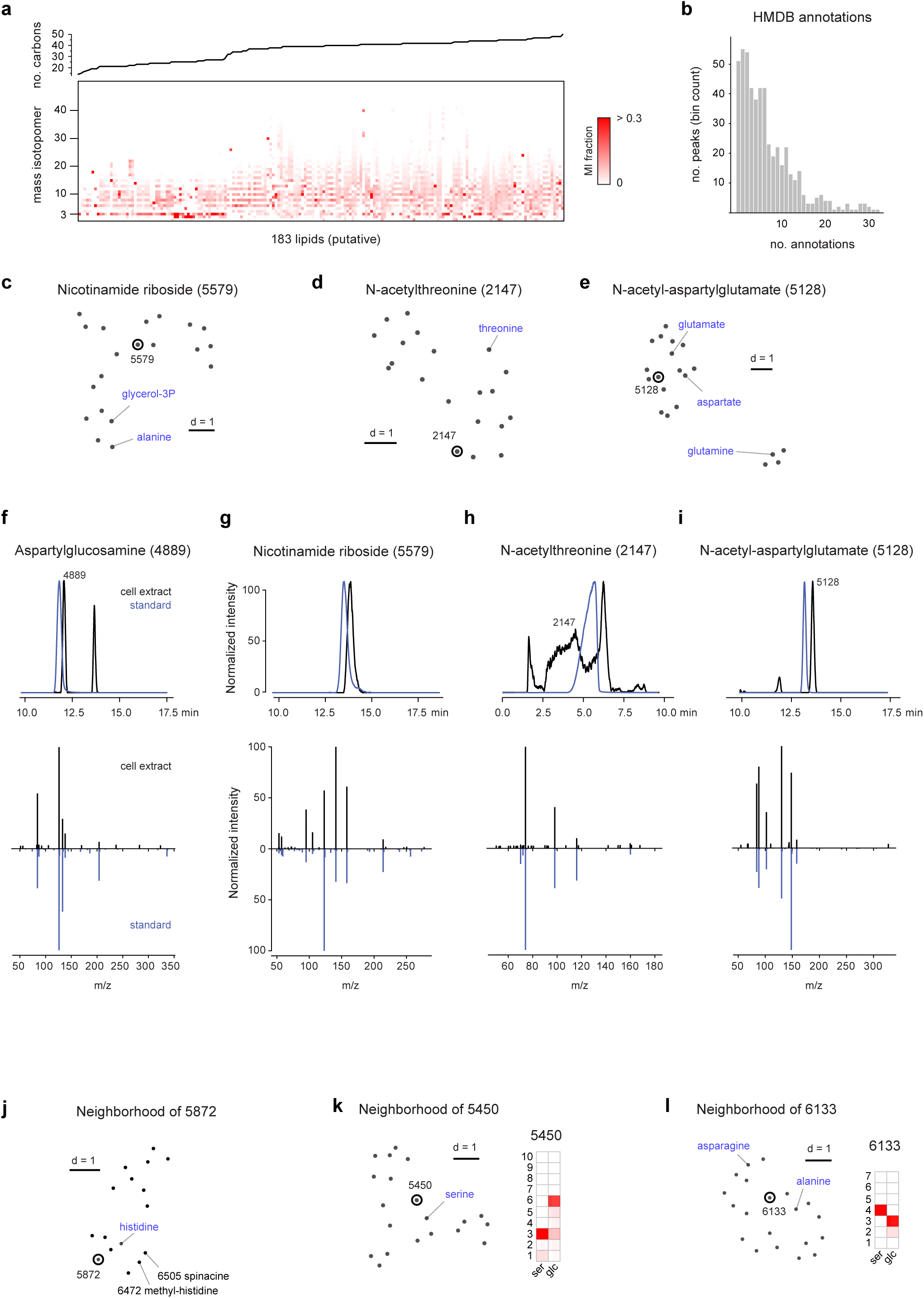
**a**, Heatmap of mass isotopomer (MI) fractions of 183 putative lipids (bottom) and their predicted number of carbons (top). **b**, Distribution of number of putative annotations per peak when matching by m/z against HMDB. **c-e**, UMap projections of neighbors of peaks 5579 (c), 2147 (d) and 5128 (e) according to the MID distance. Nearby known compounds indicated in blue. **f–g**, Validation of predicted peak identities 4889 aspartylglucosamine (f), 5579 nicotinamide riboside (g), 2147 N-acetylthreonine (h) and 5128 N-acetyl-asparatylglutamate (i) by retention time and MS^2^ fragmentation spectra of the corresponding pure standards. **j-l**, UMap projections of neighbors of peaks 5872 (j), 5450 (k) and 6133 (l) according to the MID distance, and corresponding MIDs for experiments with substantial ^13^C labeling.

**Extended Data Figure 5.**
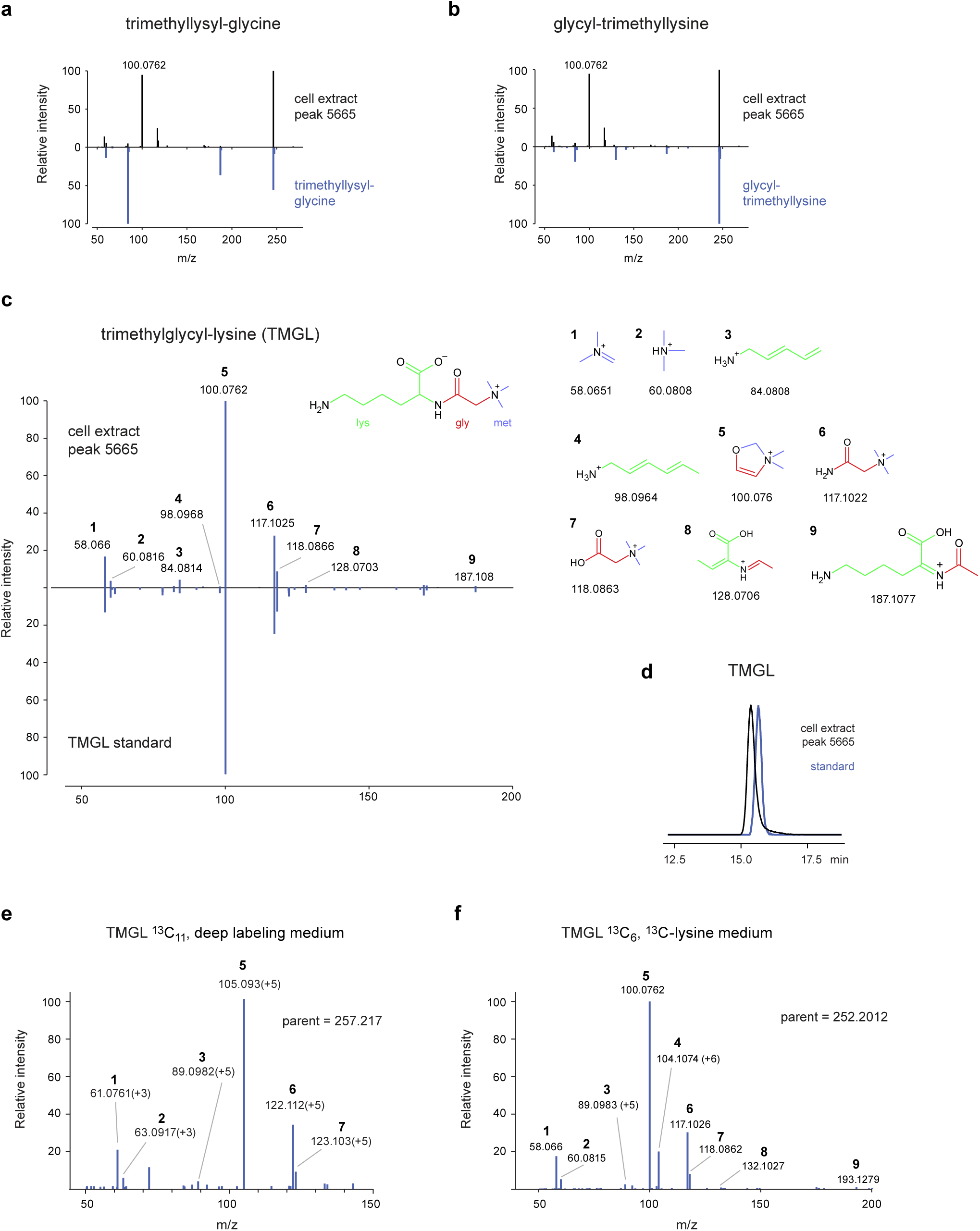
**a–b**, MS^2^ spectra of peak 5665 in cell extract and of pure standards for trimethyllysyl-glycine (a) and glycyl-trimethyllysine (b). **c**, MS^2^ spectra of peak 5665 in cell extract and of a pure trimethylglycyl-lysine (TMGL) standard. Predicted structures and theoretical m/z for indicated ions are shown, with colors indicating origin of carbons from lys, gly and met. **d**, chromatograms of peak 5665 in cell extract and a pure TMGL standard. **e**, MS^2^ spectrum of a ^13^C_11_-TMGL peak in cells cultured in “deep labeling” medium, where all amino acids and glucose are U-^13^C. Numbers refer to ion structures in (c). Mass isotopomer shifts indicated in parentheses. **f**, MS^2^ spectrum of a ^13^C_6_-TMGL peak in cells cultured in U-^13^C_6_-lysine medium, as in (e).

**Extended Data Figure 6.**
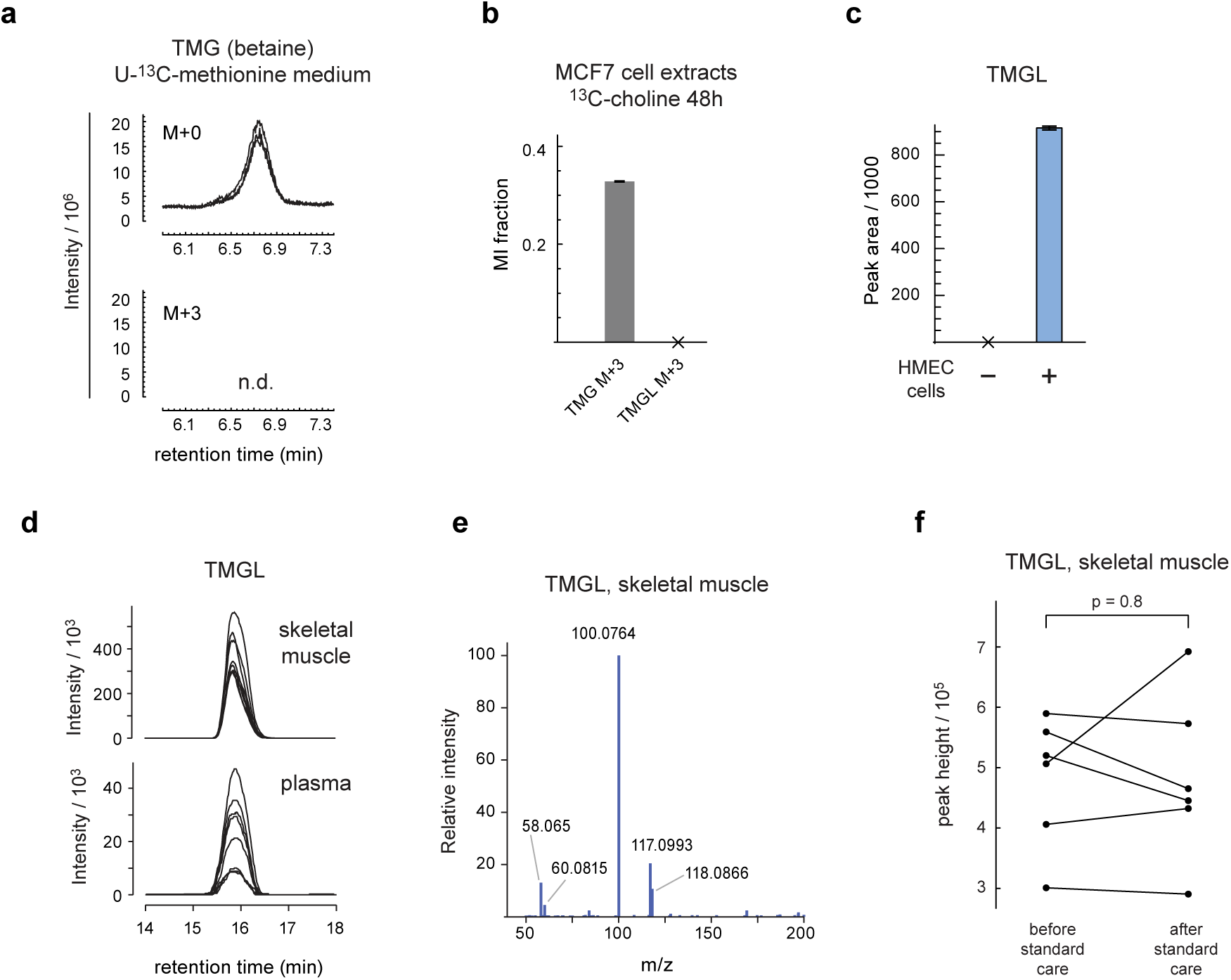
**a**, chromatograms of indicated mass isotopomers of trimethylglycine (TMG; also known as betaine) in HMECs cultured in U-^13^C-methionine medium. **b**, M+3 mass isotopomer fractions of TMG and TMGL in MCF7 cells cultured in methyl-^13^C_3_-choline medium. **c**, relative abundance of TGML in medium incubated with and without HMECs. **d**, chromatograms of TMGL from human plasma and skeletal muscle samples. **e**, MS^2^ spectrum of the TMGL peak in a human skeletal muscle sample. **f**, relative abundance of TMGL in human skeletal muscle before and after standard care (control group).

## References

1. Wang-Sattler, R., et al. Novel biomarkers for pre-diabetes identified by metabolomics. Molecular Systems Biology 8, 615 (2012).

2. Stewart, N. A., Buch, S. C., Conrads, T. P. & Branch, R. A. A UPLC-MS/MS assay of the “Pittsburgh cocktail”: six CYP probe-drug/metabolites from human plasma and urine using stable isotope dilution. Analyst 136, 605–612 (2011).

3. Buergel, T. et al. Metabolomic profiles predict individual multidisease outcomes. Nat Med 28, 2309–2320 (2022).

4. da Silva, R. R., Dorrestein, P. C. & Quinn, R. A. Illuminating the dark matter in metabolomics. Proceedings of the National Academy of Sciences 112, 12549–12550 (2015).

5. Mahieu, N. G. & Patti, G. J. Systems-Level Annotation of a Metabolomics Data Set Reduces 25 000 Features to Fewer than 1000 Unique Metabolites. Analytical Chemistry 89, 10397–10406 (2017).

6. Chen, L. et al. Metabolite discovery through global annotation of untargeted metabolomics data. Nat Methods 18, 1377–1385 (2021).

7. Xiao, J. F., Zhou, B. & Ressom, H. W. Metabolite identification and quantitation in LC-MS/MS-based metabolomics. TrAC Trends in Analytical Chemistry 32, 1–14 (2012).

8. Sumner, L. W. et al. Proposed minimum reporting standards for chemical analysis. Metabolomics 3, 211–221 (2007).

9. Alseekh, S. et al. Mass spectrometry-based metabolomics: a guide for annotation, quantification and best reporting practices. Nat Methods 18, 747–756 (2021).

10. Kanu, A. B., Dwivedi, P., Tam, M., Matz, L. & Hill Jr., H. H. Ion mobility–mass spectrometry. Journal of Mass Spectrometry 43, 1–22 (2008).

11. Horai, H. et al. MassBank: a public repository for sharing mass spectral data for life sciences. Journal of Mass Spectrometry 45, 703–714 (2010).

12. Wang, M. et al. Sharing and community curation of mass spectrometry data with Global Natural Products Social Molecular Networking. Nature Biotechnology 34, 828–837 (2016).

13. Baker, E. S. et al. METLIN-CCS: an ion mobility spectrometry collision cross section database. Nat Methods 20, 1836–1837 (2023).

14. Wishart, D. S. Computational strategies for metabolite identification in metabolomics. Bioanalysis 1, 1579–1596 (2009).

15. Djoumbou-Feunang, Y. et al. BioTransformer: a comprehensive computational tool for small molecule metabolism prediction and metabolite identification. Journal of Cheminformatics 11, 2 (2019).

16. Jeffryes, J. G. et al. MINEs: open access databases of computationally predicted enzyme promiscuity products for untargeted metabolomics. J Cheminform 7, 44 (2015).

17. Dührkop, K., Shen, H., Meusel, M., Rousu, J. & Böcker, S. Searching molecular structure databases with tandem mass spectra using CSI:FingerID. Proceedings of the National Academy of Sciences 112, 12580–12585 (2015).

18. Wolf, S., Schmidt, S., Müller-Hannemann, M. & Neumann, S. In silico fragmentation for computer assisted identification of metabolite mass spectra. BMC Bioinformatics 11, 148 (2010).

19. Watrous, J. et al. Mass spectral molecular networking of living microbial colonies. Proceedings of the National Academy of Sciences 109, E1743–E1752 (2012).

20. Nothias, L.-F. et al. Feature-based molecular networking in the GNPS analysis environment. Nat Methods 17, 905–908 (2020).

21. Hoffmann, M. A. et al. High-confidence structural annotation of metabolites absent from spectral libraries. Nat Biotechnol 40, 411–421 (2022).

22. Grankvist, N. et al. Profiling the metabolism of human cells by deep 13C labeling. Cell chemical biology 25, 1419–1427.e4 (2018).

23. Mahieu, N. G., Huang, X., Chen, Y.-J. & Patti, G. J. Credentialing Features: A Platform to Benchmark and Optimize Untargeted Metabolomic Methods. Anal. Chem. 86, 9583–9589 (2014).

24. Qiu, Y. et al. Isotopic Ratio Outlier Analysis of the S. cerevisiae Metabolome Using Accurate Mass Gas Chromatography/Time-of-Flight Mass Spectrometry: A New Method for Discovery. Analytical Chemistry 88, 2747–2754 (2016).

25. Brauman, J. I. Least Squares Analysis and Simplification of Multi-Isotope Mass Spectra. Anal. Chem. 38, 607–610 (1966).

26. Hosios, A. M. et al. Amino Acids Rather than Glucose Account for the Majority of Cell Mass in Proliferating Mammalian Cells. Developmental Cell 36, 540–549 (2016).

27. Miyake, M., Kakimoto, Y. & Sorimachi, M. Isolation and identification of β-citryl-L-glutamic acid from newborn rat brain. Biochimica et Biophysica Acta (BBA) - General Subjects 544, 656–666 (1978).

28. Nilsson, R., Roci, I., Watrous, J. & Jain, M. Estimation of flux ratios without uptake or release data: Application to serine and methionine metabolism. Metabolic Engineering 43, 137–146 (2017).

29. Jansen, R. S. et al. N-lactoyl-amino acids are ubiquitous metabolites that originate from CNDP2-mediated reverse proteolysis of lactate and amino acids. Proceedings of the National Academy of Sciences 112, 6601–6606 (2015).

30. Li, V. L. et al. An exercise-inducible metabolite that suppresses feeding and obesity. Nature 606, 785–790 (2022).

31. Roci, I., Watrous, J. D., Lagerborg, K. A., Jain, M. & Nilsson, R. Mapping choline metabolites in normal and transformed cells. Metabolomics 16, 125 (2020).

32. Yoshino, M. et al. Worksite-based intensive lifestyle therapy has profound cardiometabolic benefits in people with obesity and type 2 diabetes. Cell Metabolism 34, 1431–1441.e5 (2022).

33. Rühl, M. et al. Collisional fragmentation of central carbon metabolites in LC-MS/MS increases precision of ^13^C metabolic flux analysis. Biotechnology and bioengineering 109, 763–71 (2012).

34. Aitchison, J. The Statistical Analysis of Compositional Data. Journal of the Royal Statistical Society: Series B (Methodological) 44, 139–160 (1982).

35. Elenbaas, B. et al. Human breast cancer cells generated by oncogenic transformation of primary mammary epithelial cells. Genes & development 15, 50–65 (2001).

36. Hammond, S. L., Ham, R. G. & Stampfer, M. R. Serum-free growth of human mammary epithelial cells: rapid clonal growth in defined medium and extended serial passage with pituitary extract. Proceedings of the National Academy of Sciences of the United States of America 81, 5435–9 (1984).

37. Lyutvinskiy, Y., Watrous, J. D., Jain, M. & Nilsson, R. A Web Service Framework for Interactive Analysis of Metabolomics Data. Analytical Chemistry 89, 5713–5718 (2017).

38. Antoniewicz, M. R., Kelleher, J. K. & Stephanopoulos, G. Elementary metabolite units (EMU): a novel framework for modeling isotopic distributions. Metabolic engineering 9, 68–86 (2007).

